# Molecular mechanisms in fungal fatty acid synthase (FAS) assembly

**DOI:** 10.1101/336578

**Authors:** Manuel Fischer, Barbara Mulinacci, Mirko Joppe, Ronnald Vollrath, Kosta Konstantinidis, Peter Kötter, Luciano Ciccarelli, Janet Vonck, Dieter Oesterhelt, Martin Grininger

## Abstract

The fungal fatty acid synthase (fFAS) multienzyme is a barrel-shaped 2.6 MDa complex comprising six times eight catalytic domains. Upon barrel-formation, up to several hundred kDa large polypeptides intertwine to bury about 170,000 Å^2^ of protein surface. Functional, regulatory and structural data as well as evolutionary aspects of fFAS have been elucidated during the last decades. Notwithstanding a profound knowledge of this protein family, the biogenesis of the elaborate structure remained elusive. Remarkably, experimental data have recently demonstrated that fFAS self-assembles without the assistance of specific factors. Considering the infinitesimal probability that the barrel-shaped complex forms simply by domains approaching in the correct orientation, we were interested in understanding the sequence of events that have to orchestrate fFAS assembly. Here, we show that fFAS attains its quaternary structure along a pathway of successive domain-domain interactions, which is strongly related to the evolutionary development of this protein family. The knowledge on fFAS assembly may pave the way towards antifungal therapy, and further develops fFAS as biofactory in technological applications.

## Introduction

Fatty acid synthases (FAS) have been structurally studied during the last years, and a deep understanding about the molecular foundations of *de novo* fatty acid (FA) synthesis has been achieved ^1-3^ (**Figure S1A and B**). The architecture of fungal FAS (fFAS) was elucidated for the proteins from *Saccharomyces cerevisiae* (baker’s yeast) ^4-6^ and the thermophilic fungus *Thermomyces lanuginosus* ^7^, revealing an elaborate 2.6 MDa large α_6_β_6_ barrel-shaped complex that encapsulates fungal *de novo* FA synthesis in its interior (**Figure 1A**). The functional domains are embedded in a scaffolding matrix of multimerization and expansion elements. Acyl carrier protein (ACP) domains, shuttling substrates and intermediates inside the reaction chamber, achieve compartmentalized synthesis ^4^,^8^ (**Figure 1B and C**). The concept of metabolic crowding makes fFAS a highly efficient catalytic machinery, running synthesis at micromolar virtual concentrations of active sites and substrates ^9^. The outstanding efficacy in fungal FA synthesis is documented by (engineered) oleagenic yeast that can grow to lipid cellular contents of up to 90% ^10^. fFAS have also raised interest as biofactories in microbial production of value-added compounds from saturated carbon chains ^11-13^.

**Figure 1.**
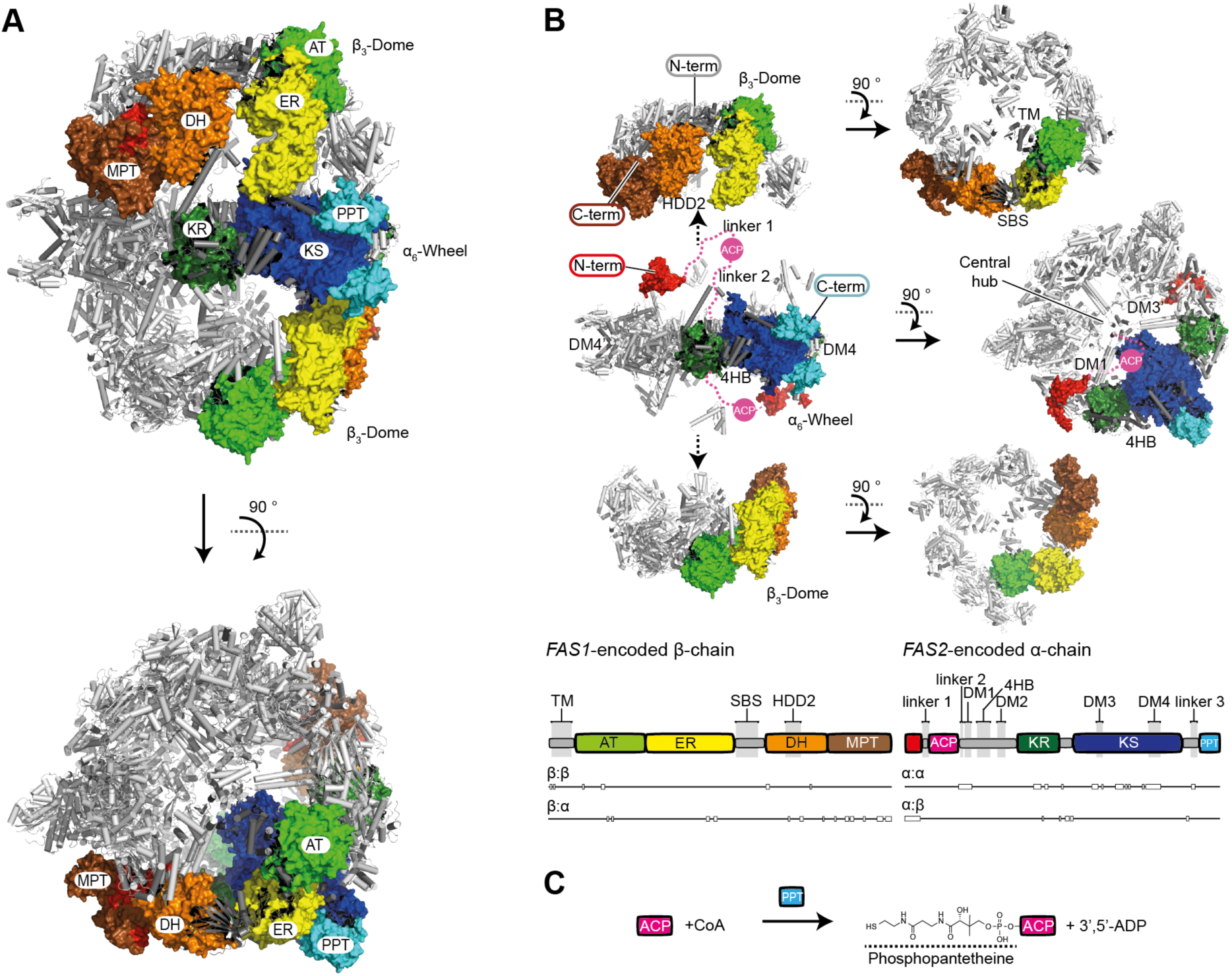
Structure of the *S. cerevisiae* FAS. A Structure of *S. cerevisiae* FAS (PDB-code: 3hmj) ^6^. Cartoon representation of the X-ray crystallographic structure shown in side (upper) and top view (lower) with two β- and two α-chains highlighted by domains in surface representation. ACP has been found in the FAS interior, but is not shown in this figure. Nomenclature: acetyl transferase (AT), enoyl reductase (ER), dehydratase (DH), malonyl-palmitoyl-transferase (MPT), acyl carrier protein (ACP), ketoacyl reductase (KR), ketoacyl synthase (KS) and phosphopantetheine transferase domain (PPT). **B** Dissection of the *S. cerevisiae* FAS barrel into the D3-symmetric α-chain hexamer (α_6_-wheel) and the two C3-symmetric β-chain trimers (β_3_-domes). β_3_-domes have been shifted for clarity (see arrows). View and coloring as in (A). ACP domains are shown for two α-chains, and are modeled by spheres in magenta. ACP linkers are indicated by dashed lines. In *S. cerevisiae* FAS, the MPT domain is distributed on both chains (β-chain part in brown and its α-chain part in red). The schematic representation of the domain architecture is attached, with the insertion elements involved in scaffolding the fFAS complex indicated. Insertion elements are highlighted in grey (nomenclature (see also above): trimerization module (TM), 6-stranded β-sheet (SBS), hotdog-domain 2 (HDD2), dimerization module 1-4 (DM1-4), 4-helical bundle (4HB)). Please note that DM2 is not visible in this structure. Rectangles attached to the domain representation indicate stretches of chain-chain interaction. **C** Scheme of the post-translational modification of ACP. For phosphopantetheinylation, ACP and PPT have to physically interact.

Facing the complexity of the fFAS structure, we recently started the project of deciphering its assembly mechanism. We were interested in two aspects. First, based on the observation that fFAS can be recombinantly expressed in *E. coli* ^14,15^, it can be posited that specific assembly factors are not required for fFAS biogenesis. Autonomous self-assembly of fFAS may essentially be envisioned by distributing the complexity of the assembly process onto a sequence of domain-domain interactions that are formed one after another. We aimed to explore this sequence of events and to analyze whether it can be correlated to the evolutionary development of fFAS, since it has been suggested that assembly pathways generally reflect protein evolution ^16^. Second, we sought to evaluate whether the knowledge on fFAS assembly may be exploited for inhibiting *de novo* fungal FA synthesis in selective antifungal therapy ^17-19^, as well as for designing fFAS based biofactories ^20^.

Our studies of the *S. cerevisiae* FAS assembly were greatly aided by engineering fFAS on the basis of the available atomic resolution models ^4-7,21^. Wildtype and several engineered *S. cerevisiae* FAS constructs were used for complementing a FAS-deficient yeast strain. Full-length and truncated *S. cerevisiae* FAS constructs were further recombinantly expressed in *Escherichia coli*. These tools in hand, we were able to address fFAS assembly in a “forward-approach”, which means that instead of often-performed dissociation based (“reverse”) approaches, we generated information based on halted assembly states and truncated structures. Here, we present a multitude of data suggesting that *S. cerevisiae* FAS autonomously assembles via a single dominant pathway, which we outline in three key processes and correlate to the evolutionary development of fFAS. The molecular details of fFAS biogenesis provide the basis for a structure-based design of assembly inhibitors and may further pave the way for designing complex compartmentalized synthetic pathways.

## Results

### The fFAS family is topologically heterogeneous on gene level

Genome sequence analysis has characterized fFAS as a heterogeneous family comprising different gene-topological variants (**Figure 2A**). As most evident gene-topological variation, fFAS are either encoded by single genes or by two genes. Two-gene-encoded fFAS appear to originate from a single-gene encoded precursor split into two at various fission sites that are generally located within domains ^3,22^. In *S. cerevisiae* and *T. lanuginosus* FAS, both representing the *Ascomycota*-type fFAS, the C-terminus of the β-chain and the N-terminus of the α-chain intertwine to form the MPT domain (**Figure 2Bi**) ^4,7^. In *Tremellomycetes*-type fFAS, the termini of polypeptide chains form a 4-helical bundle (4HB) at the interface of the KR and the KS domain (**Figure 2Bii**). At the *Rhodosporidium toruloides* FAS fission site chains share an antiparallel β-sheet (SBS) domain, but, different to *Ascomycota*- and *Tremellomycetes*-type FAS, the termini do not intertwine ^14^ (**Figure 2Biii**). Gene topological variations are also apparent in the distribution of insertion elements that scaffold the fFAS barrel (see **Figure 1** and **2A**). While the dimerization module DM3 is highly conserved in type I FAS ^22^, the trimerization module (TM) and dimerization module (DM2) do not occur in ancestral variants and the evolutionarily related bacterial type I FAS.

**Figure 2.**
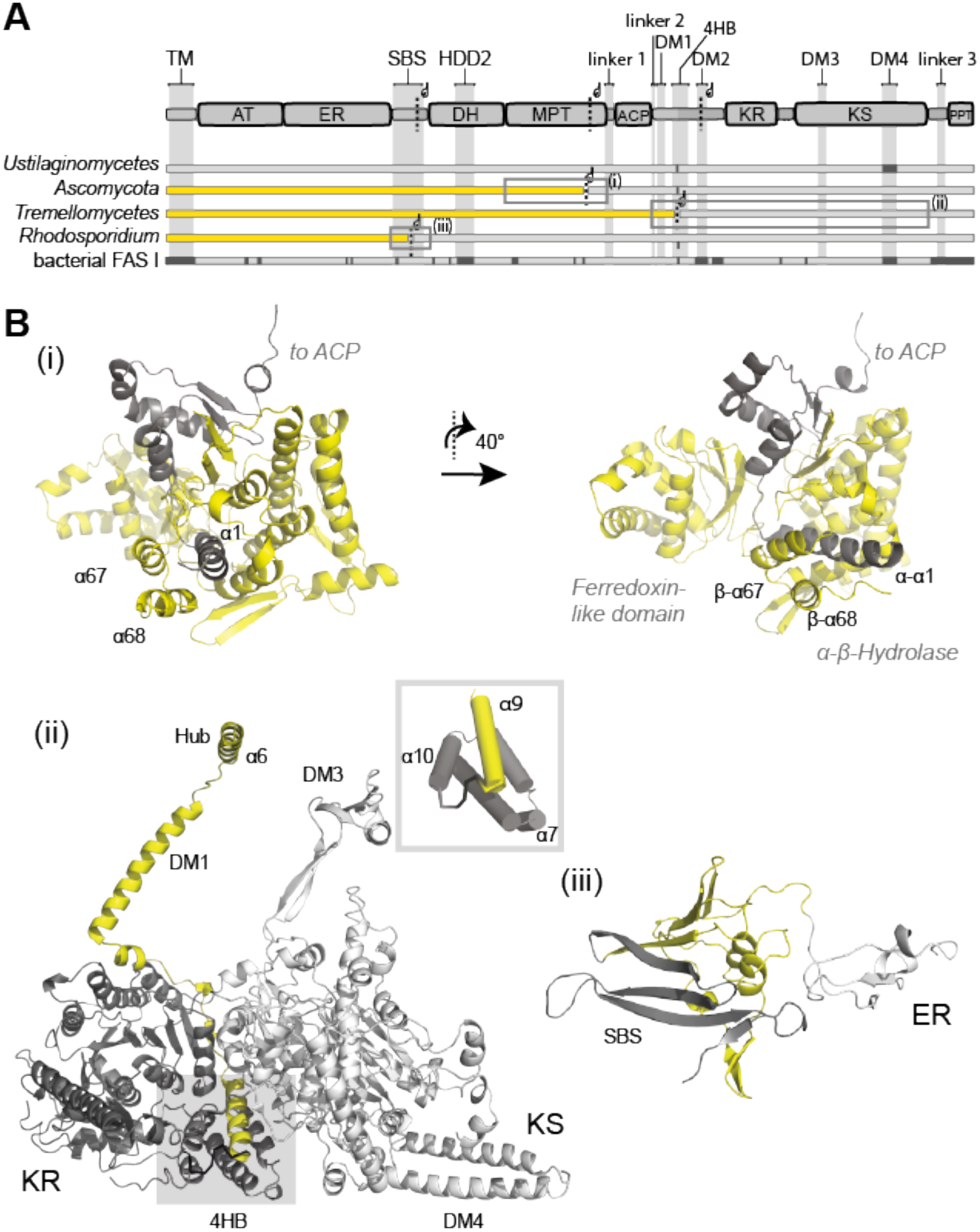
The fFAS family. A Topological variants of fFAS. Domain architecture is given for single-chain fFAS. Abbreviations used as in **Figure 1**. Four fFAS variants differing in fission sites as well as in the distribution of insertion elements are given (missing insertions in dark grey). *Ustilaginomycetes*-type FAS carries all domains on a single chain (single polypeptide), *Ascomycetes*-type (including *S. cerevisiae* and *C. albicans*), *Tremellomycetes*- type (including *C. neoformans* and *C. gattii*) and *Rhodosporidium*-type FAS I are two-gene encoded variants. Substructures of *S. cerevisiae* FAS shown in panel B are highlighted by grey frames. Yellow and grey coloring indicate *FAS1*-encoded polypeptides (β-chains) and *FAS2*-encoded polypeptides (α-chains), respectively. **B** Substructures of *S. cerevisiae* FAS depicting the fission sites in fFAS variants. Secondary structure elements are shown in *S. cerevisiae* FAS numbering as introduced by Jenni *et al.* ^7^. Coloring as in **Figure 2A**. Fission site of *Ascomycetes*-type FAS within the MPT domain is shown in two orientations **(i)**, of *Tremellomycetes*-type FAS within the 4HB (see also inset) (ii), and of *Rhodosporidium*-type FAS with the antiparallel β-sheet domain (SBS) (iii). For an extended version of this figure see **Figure S2**.

In a first experiment, we analyzed the assembly of above described fFAS variants by constructing gene topologies with *S. cerevisiae* FAS. The fFAS variants were rebuilt in *S. cerevisiae* FAS by initially engineering a single-gene encoding fFAS with *FAS1* and *FAS2* connected by a sequence that natively links the two genes in *Ustilago maydis* FAS (*Sc_fas1-fas2*). Taking this construct as a template, we then engineered splitting sites as occurring in *Tremellomycetes*-type (*Sc_Tre*) and *Rhodosporidium*- type FAS (*Sc_Rho*) (see **Figure 2Bii** and **iii,** and **Figure S2A** and **B**). In the experimental procedure, a FAS-deficient *S. cerevisiae* strain, growing on external FA, was complemented by plasmids encoding the fFAS variants (**Table S1**) ^23,24^. All three constructs successfully complemented the deficiency in *de novo* FA synthesis of the FAS-deficient yeast, as read-out by growth rates in FA-limited liquid cultures and by spot dilutions on medium without added FA. We also performed Native-PAGE with Western-Blot detection to visualize intact barrel-shaped FAS in *S. cerevisiae* cytosolic fractions. For analysis, we blotted cell lysates of the complemented FAS-deficient *S. cerevisiae* strains separated by Native-PAGE, and made *S. cerevisiae* FAS visible with polyclonal rabbit anti-FAS antibodies ^25^. All three constructs successfully complemented the deficiency in *de novo* FA synthesis of the FAS-deficient yeast, as well as assembled to the barrel-shaped complex (**Figure S3A-C**). Our data show that the fission event, which splits the single-gene encoding fFAS into two-gene encoding fFAS and is a late step in evolution ^3,22^, is rebuild early in the assembly by the interaction of chain termini. This early interaction may also be seen as event happening prior to the actual assembly (as the specific process of barrel formation) that captures all variants to assemble via a single assembly pathway; in line with the conception of the high evolutionary conservation of assembly pathways in protein families ^26^. We term this assembly step “pseudo-single chain formation” in the following.

To evaluate the impact of insertion elements on fFAS assembly, we further engineered fFAS constructs with the β-chain’s N-terminal trimerization module TM or/and the α-chain embedded dimerization module DM2 deleted (see **Figure 2A**). The TM closes the barrel at its apical sites and DM2 is placed at the outer barrel perimeter. DM2 increases the KR/KR interface as shown in the crystal structure of *T. lanuginosus* FAS ^7^. Native PAGE Western Blot analysis indicates intact assembly of the deletion mutants, again demonstrating the high evolutionary conservation of the assembly pathway within the fFAS family (**Figure 3A**). Further protein properties were determined *in vitro* on the purified proteins by performing size exclusion chromatography (SEC), a thermal shift assay (TSA) and an enzymatic activity assay with the purified Strep-tagged constructs (**Figure 3B-C**). SEC and TSA data show compromised stability of the proteins with deleted insertion elements, documented by an increased tendency to aggregation in SEC and a drop in protein melting temperature in TSA. A decrease in overall FA synthesizing activity from initial 2953 ± 205 mU/mg for wildtype FAS to 833 ± 73 mU/mg for the ΔTM deletion, 1069 ± 97 mU/mg for the ΔDM2 deletion and to 466 ± 125 mU/mg for the double deletion well correlates with protein stability measures. Data indicate that the insertion modules TM and DM2 are not essential for assembly, but stabilize the fFAS barrel and increase the fFAS catalytic efficacy. Many of the scaffolding elements, i.e. the elements that are not strictly conserved within the fFAS protein family like TM and DM2, have appeared at a later stage during protein evolution, when the elaborate barrel-shaped fold had been already developed ^22^.

**Figure 3.**
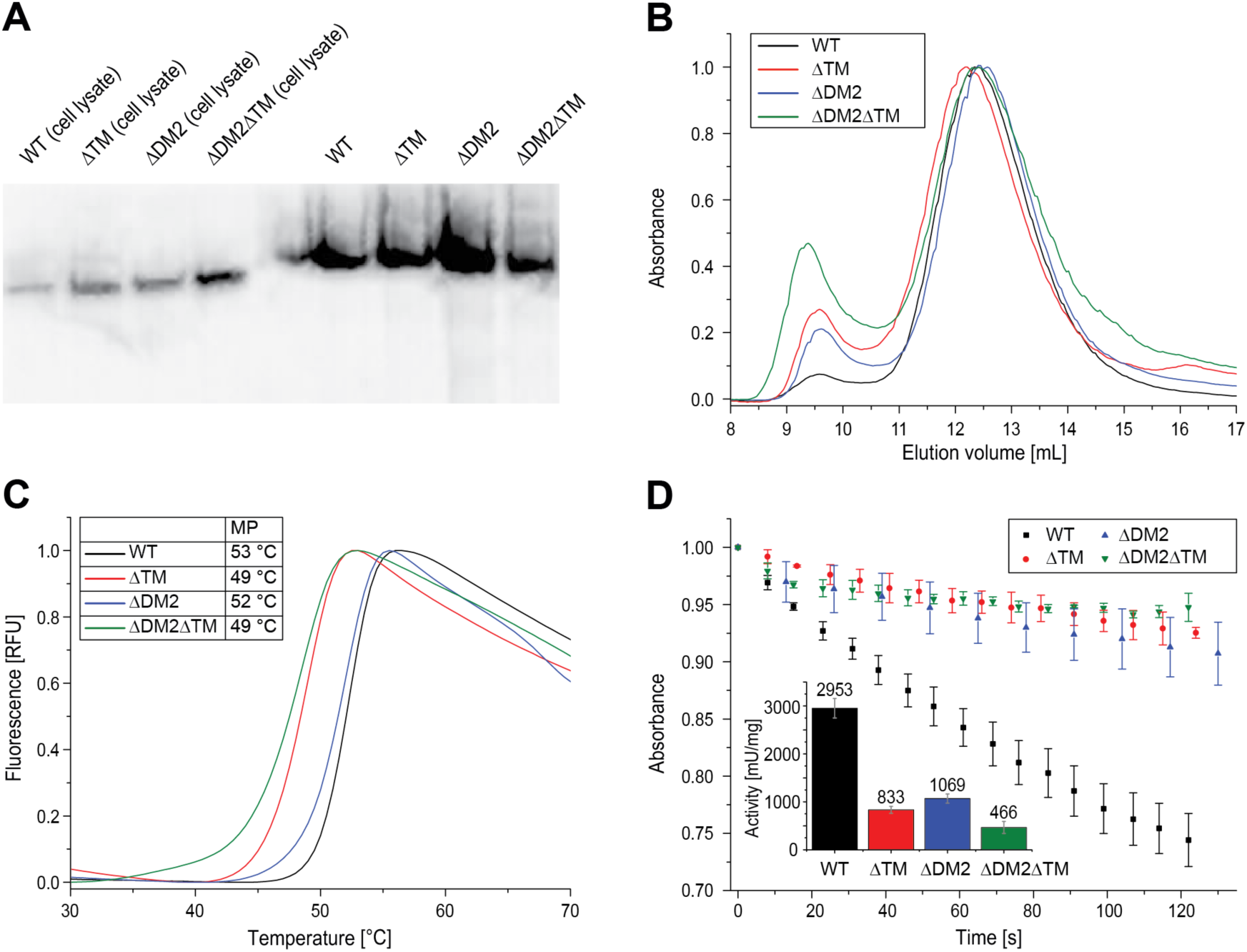
Purification and analysis of fFAS with deletion of insertion elements. A Native PAGE Western Blot of *S. cerevisiae* FAS constructs from cell lysates (left) and after purification (right) as indicated. WT = wild type. **B** FAS constructs as shown in panel A were purified with SEC (Superose 6 increase 10/300 GL, buffer: 100 mM sodium phosphate pH = 6.5, 200 mM sodium chloride). UV absorption at 280 nm has been normalized. **C** Typical melting curves of FAS constructs received in TSA. Fluorescence has been normalized. Average melting temperatures (MP) of two technical replicates are shown in inset table. The difference between technical replicates was smaller than 0.5 °C. **D** Activity assay of FAS constructs shows as time course of NADPH absorption at 334 nm. Inset diagram shows calculated specific activities (in mU/mg) as bars. Average and standard deviation (±1 σ) of three technical replicates are shown for each construct.

### “Pseudo-single chain” formation is an early event in fFAS assembly

To validate the relevance of pseudo-single chain formation as an early step in fFAS assembly, we generated a set of *S. cerevisiae* FAS mutants that modulate the MPT interface. The MPT interface contributes a marginal amount to the overall about 170,000 Å^2^ of protein surface being buried upon barrel formation ^7^. Therefore, we assumed that MPT mutants should only affect barrel assembly, if MPT formation by the α- and the β-chain indeed constitutes an early event in the assembly. We initially tested two FAS constructs; (i) β-chain deleted in the C-terminal helices α67 and α68 (pRS415_*fas1Δ*α67/68) combined with wildtype α-chain (pRS413_*FAS2*) yielding strain *Sc_Δα67/68*, and (ii) wildtype β-chain (pRS415_*FAS1*) combined with α-chain deleted in about half of the N-terminal α1-helix (amino acids K2 to H11; pRS413_*fas2Δα1(2-11)*) giving strain *Sc_Δα1(2-11)*. Both complementation constructs reduce the interface of α-chain/β-chain interactions in the MPT domain. Both of these constructs did not or just very poorly restore *de novo* FA synthesis in the FAS-deficient yeast strain (**Figure 4A** and **B**). As the complementation assay does not indicate whether absent activity can indeed be attributed to an abolished assembly or is rather caused by the compromised catalytic activity of assembled fFAS, we also performed Native-PAGE with Western-Blot detection. Both strains *Sc_Δα67/68* and *Sc_Δα1(2-11)* did not contain assembled FAS (**Figure 4C**). Non-assembled polypeptides were not visible in Native-PAGE, which is best interpreted as degradation of non-assembled FAS as suggested earlier ^25,27^. Absence of assembled FAS, as a result of transcriptional down-regulation or RNA instability instead of assembly failure, can be excluded for the *fas2Δαas2-11)-* construct based on a previous study showing constant expression of a *fas2-lacZ*-fusion-gene lacking the first 39 nucleotides of the *FAS2* open reading frame ^28^.

**Figure 4.**
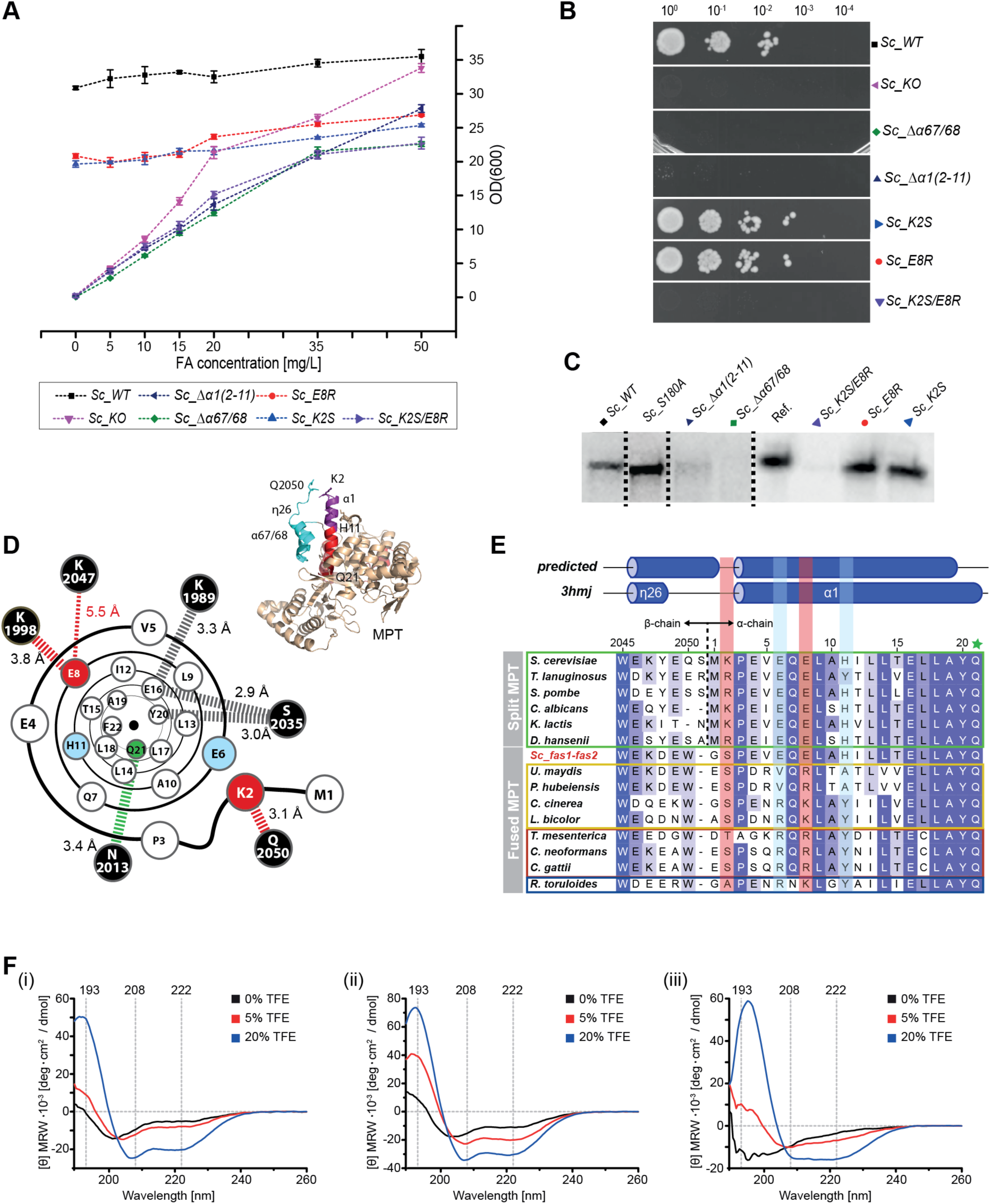
Interaction of α-chain and β-chain in the MPT domain for pseudo-single chain formation. Details and abbreviations of complementation constructs are outlined in **Table 1**. **A** Growth behavior of mutated strains in liquid cultures supplemented with external FA. Experimentally determined values (each in 5 technical replicates; error bars represent ±3 σ); measuring points are connected by dashed lines for clarity. For biological replicates, see **Figure S4A-C** and **Table S2**. Please note that the relatively higher ODs for WT and KO originate from deviant starting conditions, as they were precultured in YPD-FA instead of SD-FA medium (see Supplemental Information). **B** Ten-fold serial dilutions (starting from OD(600) = 1) of log-phase cultures spotted on YPD agar without external FA supply after incubation for 48 h at 30 °C. **C** Native-PAGE-Western-Blot analysis of FAS from mutant strains grown to the log-phase. Bands indicate presence or absence of intact FAS barrels. As reference, we used purified FAS from *S. cerevisiae*. For clarity, the figure has been assembled from different blots as indicated by dashed lines. For the complete blots, see **Figure S5A-C**. **D** Cartoon illustrating the α1-helix with key polar interactions to the β-chain (black spheres) represented by dashed bars. Distances as indicated are calculated from the X-ray structure (PDB-code: 3hmj) ^21^. Amino acids that were sensitive to mutations in the FAS assembly process are shown as red spheres; insensitive mutations are shown as blue spheres, and the catalytic Q21 (involved in the catalytic triad of the MPT active site) as green sphere. For guidance, the MPT domain of *S. cerevisiae FAS* is shown as inset. **E** Alignment of sequences covering the α1-helix (*S. cerevisiae* FAS numbering). Sequences include *Ascomycota*-type FAS (green box), single-gene encoded fFAS (yellow), *Tremellomycetes*-type FAS (red), *Rhodosporidium*-type FAS (blue) and the engineered *fas1-fas2*-fusion strain *Sc-fas1-fas2*. The alignment was created with Clustal Omega on the EBI webserver based on the full length FAS sequence and colored according to occurrence ^30^. Two-genes encoded FAS were submitted as *FAS1*-*FAS2*- fusions. Predicted *S. cerevisiae* FAS secondary structure from PsiPred ^31^ and the secondary structure as observed in the X-ray crystal structure (PDB-code: 3hmj) are attached. Loci that are mutation sensitive in *Ascomycota*-type FAS assembly are highlighted by a red background; two further loci, which we have exchanged in mutational studies are in blue, and the catalytically relevant Q21 is indicated by a green star. Uniprot (or GenBank in case of *Tremella mesenterica*) accession numbers of sequences are: *Candida albicans* (P34731, P43098), *Coprinopsis cinerea* (A8NUB3), *Cryptococcus gattii* (E6R622, E6R621), *Cryptococcus neoformans* (Q5KG98, Q5KG99), *Debaryomyces hansenii* (Q6BWV8, Q6BWN1), *Kluyveromyces lactis* (Q6CWN6, Q6CT25), *Laccaria bicolor* (B0D9Q1), *Pseudozyma hubeiensis* (R9P8H2), *Rhodosporidium toruloides* (M7WSW5, M7XM89), *Saccharomyces cerevisiae* (P07149, P19097), *Schizzosaccharomyces pombe* (Q9UUG0, Q10289), *Thermomyces lanuginosus* (A4VCJ6, A4VCJ7), *Tremella mesenterica* (XP_007006732.1, XP_007006745.1), *Ustilago maydis* (A0A0D1C5S0). **F** Analysis of custom-synthesized peptide fragments CD-spectroscopy recorded at different TFE concentrations; peptides α1 (**i**), K2S-E8R-mutated α1 (**ii**) and in α67/68 (**iii**). For growth behavior of additional mutants, see **Figure S4C, Tables S3**, **S4 and S5).**

We further modulated the interface of chains in the two-gene encoding fFAS variants *Sc_Tre* and *Sc_Rho* (see **Figure S3A-C**). We deleted the N-terminal three α-helices of the *Tremellomycetes*-type mimicking FAS α-chain and the N-terminal β-sheet of the *Rhodosporidium*-type mimicking FAS α- chain giving strains *Sc_Tre_Δα10-12* and *Sc_Rho_Δ(1-53)*, respectively. Both strains failed to restore *de novo* FA synthesis, demonstrating the broad impact of the early interaction of termini (pseudo-single chain formation) in fFAS assembly (**Figure S4A-C**).

**Table 1.** *S. cerevisiae* FAS strains used in this study. The ability for complementation is shown in **Figure 4A-C** and **Figure S4A**.

| Strain | FAS genotype (on pRS) |
| --- | --- |
| *Sc* KO | $\Delta fas1 / \Delta fas2$ |
| *Sc* WT | *FAS1 / FAS2* |
| *Sc* K2S | *FAS1 / fas2_K2S* |
| *Sc* E8R | *FAS1 / fas2_E8R* |
| *Sc* K2S/E8R | *FAS1 / fas2_K2S/E8R* |
| *Sc* $\Delta\alpha$1(2-11) | *FAS1 / fas2_Δα1(2-11)* |
| *Sc* $\Delta\alpha$67/68 | *fas1_Δα67/68 / FAS2* |
| *Sc* S180A | *FAS1 / fas2_S180A* |

As a next step in the analysis of the chain interaction, we mutated the MPT interface of *S. cerevisiae* FAS and cloned constructs with point mutations in helix α1 (**Figure 4D)**. Amino acids K2, E6, E8 and H11 were selected as candidates based on their conservation in *Ascomycota*-type FAS, and mutated to their most frequent exchanges in non-*Ascomycota*-type FAS. (**Figure 4E**). All single mutated constructs (pRS415_*FAS1*; mutated pRS413_*fas2\**; K2S, E6V, E8R and H11A) were able to restore *de novo* FA synthesis in the FAS deficient yeast strain. However, double mutated constructs, permutating the above amino acid exchanges, identified K2S-E8R-double mutated FAS as assembly deficient (see **Figure 4A-C**). To better understand the impact of mutations, we analyzed custom-synthesized peptide fragments in their secondary structure by CD-spectroscopy in co-solvents ^29^. We observed a high propensity of α1-peptide to form a helix, which was even more pronounced in the K2S-E8R-mutated peptide (**Figure 4F**). According to these data, the assembly defect by the K2S-E8R mutation, as observed in the complementation assay (see **Figure 4A-E**), may either originate from changed specific interaction by amino acid exchanges, or from changed α-helical properties of the mutated α1-helix that interferes in local assembly properties.

### Post-translational modification occurs within a dimeric sub-structure

In a stepwise deconstruction approach, we dissected fFAS into domains and multi-domain constructs, which we then analyzed in structural properties and catalytic activity. Since the proteolytic degradation of the *S. cerevisiae* FAS chains has been reported as a regulatory step of α/β expression ^25^, we expressed proteins recombinantly in *E. coli*. We demonstrated the suitability of *E. coli* as an expression host by successfully producing *S. cerevisiae* FAS and an *Ustilaginomycetes*-type mimicking *fas1*-*fas2* fusion protein. This was not unexpected, since the expression of other fFAS constructs in *E. coli* as well as the fFAS homologous bacterial type I FAS occurring in *Corynebacteria, Mycobacteria* and *Nocardia* (CMN-bacterial FAS) has been reported before (**Figure S7A-D**) ^14,15,32,33^. For the deconstruction approach, we focused on the α-chain. The α-chain harbors the relevant domains for post-translational modification as well as the domains contributing most of the overall 170,000 Å^2^ buried surface of fFAS. The β-chain was not proteolytically stable as a separate protein in *E. coli*, which impaired *in vitro* assembly experiments.

We dissected the α-chain from its C-terminus, and initially probed the role of the C-terminal PPT as a separate domain and as part of larger constructs. The structural frame of the post-translational phosphopantetheinylation can provide an important snapshot in fFAS assembly. The phosphopantetheinylation reaction in fFAS is largely restrained by the requirements of the domains ACP and PPT. As has been reported before, the ACP is monomeric, whereas the PPT domain is only active in a multimeric state ^21^. As a separate domain, PPT occurs as a trimer ^21^, as also described for the bacterial homolog AcpS ^34^. SEC analysis of a KS-PPT di-domain construct showed a dimeric character of the KS-PPT substructure. It seems that the large dimeric interface of the KS dimer overrides the PPT trimeric preference. Constructs PPT and KS-PPT as well as *S. cerevisiae* FAS were phosphopantetheinylation-active (see **Figure 5A** and **Figure S8A**), indicating that the PPT domain is active in both its dimeric and trimeric state ^21^. While ACP and PPT have to physically interact during phosphopantetheinylation, we were not able to identify stable ACP:PPT complexes by pull-down, co-purification and crosslinking experiments, indicating that the interaction is transient and unstable (**Figure S6A-C**).

**Figure 5.**
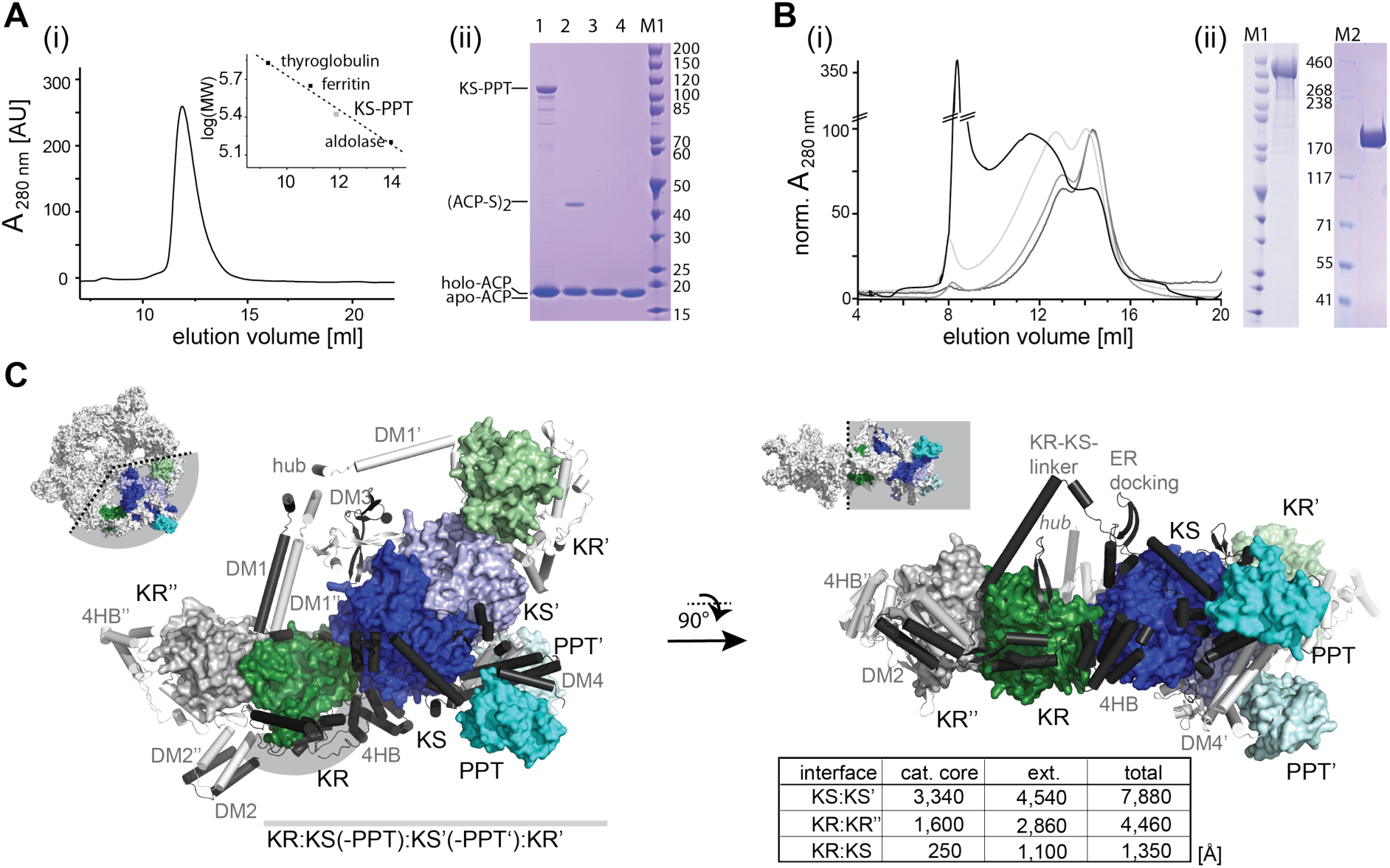
Oligomeric requirements of α-chain domains. A Analysis of the KS-PPT di-domain construct. (i) SEC on a Superdex 200 Tricorn 30/100 column and calibration curve in inset. The peak corresponds to an apparent molecular weight of 250 kDa, equivalent to a stoichiometry of 2.5 (calculated molecular weight 99.5 kDa; see also **Figure S6**), suggesting a dimer with increased apparent weight possibly due to a non-globular shape of the protein. (ii) 4-12% Bis/Tris SDS-PAGE gel (NuPage, Invitrogen) of a phosphopantetheinylation assay. Holo-ACP tends to dimerize *via* a disulfide formation, giving (ACP-S)_2_, which can be used as read-out for PPT activity. The disulfide bond is cleaved under the reducing conditions of the sample loading buffer. Lane M, marker; 1, reaction solution; 2, ACP purified from the reaction solution and loaded on gel under non-reducing conditions; 3, same as 2 but loaded on gel under reducing conditions; 4, apo-ACP reference. For an extended presentation of data see **Figure S8A**. **B** Analysis of the αΔMPT-ACP construct. SEC on a Superose 6 Tricorn 30/100 column with protein preparations from purifications under native and denaturing conditions. Peaks correspond to apparent molecular weights of 400 kDa (○), 600 kDa (Δ) and 800 kDa (□). The sharp peak in at about 8 ml corresponds to protein aggregates eluting in the void volume. (ii) SDS-PAGE gels of purified αΔMPT-ACP (see also **Figure S8B**). **C** KR:KS(-PPT):KS’(-PPT’):KR’ substructure and analysis of interfaces. Catalytic cores are colored as introduced in **Figure 1**. Insertions are shown in cartoon representation in black. Interfaces are listed as table, and numbers are given for the catalytic cores (cat. core) and the contributions by insertion elements (ext.). For stabilizing the KR:KS interface, a large insertion, including the DM1-4HB connecting linker and 4HB, enwrap the KR (insertion highlighted by grey background). For orientation, α6-wheel substructures are shown in in insets. Calculation of interfaces and their representation in this figure are based on *S. cerevisiae* FAS data ^21^ with modeled DM2 ^7^.

N-terminal elongation of the KS-PPT construct by a sequence including DM1, DM2 and KR (construct termed α_ΔMPT-ACP) led to protein aggregation (**Figure 5B**). SEC analysis resulted in a sharp peak at an apparent mass of approximately 450 kDa, indicating a fraction of dimeric species, but mainly showed unspecific higher oligomerization/aggregation. Data do not support the formation of stable α_6_- wheel structures formed by α_ΔMPT-ACP. SEC elution fractions of α_ΔMPT-ACP still showed PPT-activity, which implies that structured dimeric KS-PPT cores remain intact upon aggregation (**Figure S8B)**. Higher ratios of dimeric species were received, when the protein was purified under denaturing conditions and refolded by SEC or dialysis under low protein concentrations (see **Figure 5B**). Further N-terminal elongation to a α_ΔMPT construct as well as to the full-length α-chain did not resolve aggregation formation. Intriguingly, in spite of aggregation, the ACP domain of α_ΔMPT was quantitatively phosphopantetheinylated (*in cis*). Phosphopantetheinylation of ACP was probed by inserting a TEV-proteolytic cleavage site in the linker C-terminal to ACP, allowing ESI-MS analysis of the separate ACP received after TEV-proteolytic digestion of α_ΔMPT aggregates (**Figure S9**).

Data collected on the truncated α-chain constructs imply that the phosphopantetheinylation active species is dimeric, organized by the KS dimer as the prominent structural unit. It can further be concluded that the sequence ACP-KR-KS-PPT bears the information for forming the phosphopantetheinylation competent complex, but not for forming the D3 symmetric α_6_-wheel structures. Since the α-chain constructs run into aggregation, but are nevertheless phosphopantetheinylated, it seems that the phosphopantetheinylation status of fFAS is not proofread during assembly. For confirming this result, we analyzed the phosphopantetheinylation-deficient S180A *S. cerevisiae* FAS in our assembly assay (see **Figure 3C**). The mutated construct was unable to restore *de novo* FA synthetic activity in the complementation assay, but indeed assembled to the α_6_β_6_ complex, supporting an assembly process that does not supervise post-translational phosphopantetheinylation. This observation is in agreement with a similar result received earlier ^35^.

### Pseudo-single chain formation may be targeted in antifungal therapy

Fungal infections are an emerging threat to mankind ^36^. Human pathogens are, among others, *Cryptococcus neoformans, Cryptococcus gattii* and species of the genus *Candida* ^37^. *C. neoformans* and *C. gattii* are the causative agents of cryptococcosis, responsible for cutaneous and pulmonary infections, as well as meningitis ^38^. *Candida* species are the most important causes of opportunistic mycosis, and responsible for mucosal, cutaneous and also invasive infections ^39^. Instead of targeting active sites, which are conserved throughout bacteria and eukaryotes, assembly inhibition holds out the prospect for an ultra-selective antifungal therapy that leaves mammalian FAS I system as well as the mitochondrial and bacterial type II systems unaffected ^3^. The early event of pseudo-single chain formation of fFAS may be of foremost relevance in such a concept, as it provides the chance to impair assembly by targeting a comparable small interface. For testing the approach, we designed constructs, in which the β-chain is elongated for internal competition in pseudo-single chain formation. The elongation of the 11 amino acid long α1-helix already compromised the ability to restore FA *de novo* synthesis, and the fusion of complete part α-chain coded MPT turned out to be lethal (see **Figure S4A-C**).

## Discussion

Data presented in this study indicate that the assembly of the α_6_β_6_ *S. cerevisiae* FAS essentially organizes into three key processes (**Figure 6**): (1) The collaboration of α- and β-chain termini for the formation of pseudo-single chains is an early event in *S. cerevisiae* FAS assembly. The α-chain N-terminus, likely already developed in its secondary structure, intertwines with a β-chain C-terminus by getting sandwiched between a structured MPT core fold and a α67/α68 element that may fold only upon interaction ^40^. This initial interaction is sensitive to perturbations as indicated by two experimental set-ups. First, site directed mutagenesis identified two residues at the interface that are crucial for assembly and highly conserved in *Ascomycota*-type FAS (see **Figure 2A-F**). Second, the initial interaction can be competitively inhibited *in cis* (see **Figure S4 A-C**). This interaction may be an interesting target for antifungal therapy. (2) The second step in assembly is the formation of a dimeric unit as a platform for post-translational phosphopantetheinylation by the interaction of the domains ACP and PPT. The *S. cerevisiae* FAS KS:KS interface is the largest interface, comprising 7,880 Å^2^, and was characterized as evolutionarily ancient ^22^. Similarly as having recruited other domains by gene fusion during the course of evolution, the KS dimer appears to have conserved its central role as nucleation site for assembly ^16,26^. As disclosed by recombinant expressions in *E. coli* (see **Figure 5A-C**, and **Figure S8A** and **B**), the dominant structural role of the KS dimer is also evident in *S. cerevisiae* FAS subconstructs KS-PPT and α_*Δ* MPT-ACP (domains structure KR-KS-PPT). (3) The barrel-shaped structure encloses in a third and last process of assembly. On the way to forming the mature α_6_β_6_-complex, KR dimerization (including the adjacent scaffolding domains DM1 and DM2) contributes the largest remaining interface of 4,660 Å^2^ (13,480 Å^2^ upon α_6_-wheel assembly, see **Figure 5C**). The β-chain adds comparably small interfaces. Here, most relevant is the AT domain interacting with the domains ER and MPT of the neighboring chains, as well as the trimerization of the TM scaffolding elements (see **Figure 1B**). fFAS specific insertions elements, as e.g. DM2 as well as the trimerization domain TM, contribute to barrel stability, but are not of crucial importance for barrel biogenesis (see **Figure 5A-C**). This is also likely true for α-β connecting insertion elements (termed C-connections) that mediate interactions between α_6_-wheel and β_3_-domes. The C1 insertion of the ER-domain mediating the interaction with the KS, the C2 hotdog domain insertion of DH-domain mediating the interaction with the KR-KS connecting helix, and the C3 insertion mediating the docking of the MPT-domain to the KR’ are not uniformly distributed within the fFAS family (including ancestral fFAS and CMN-bacterial FAS) ^33,41^, which speaks against their crucial function in fFAS assembly. Rather these the C-connections stabilize the barrel for improved protein stability and/or increased catalytic efficiency, as demonstrated experimentally for insertion elements DM2 and TM (see **Figure 3A-C**).

**Figure 6.**
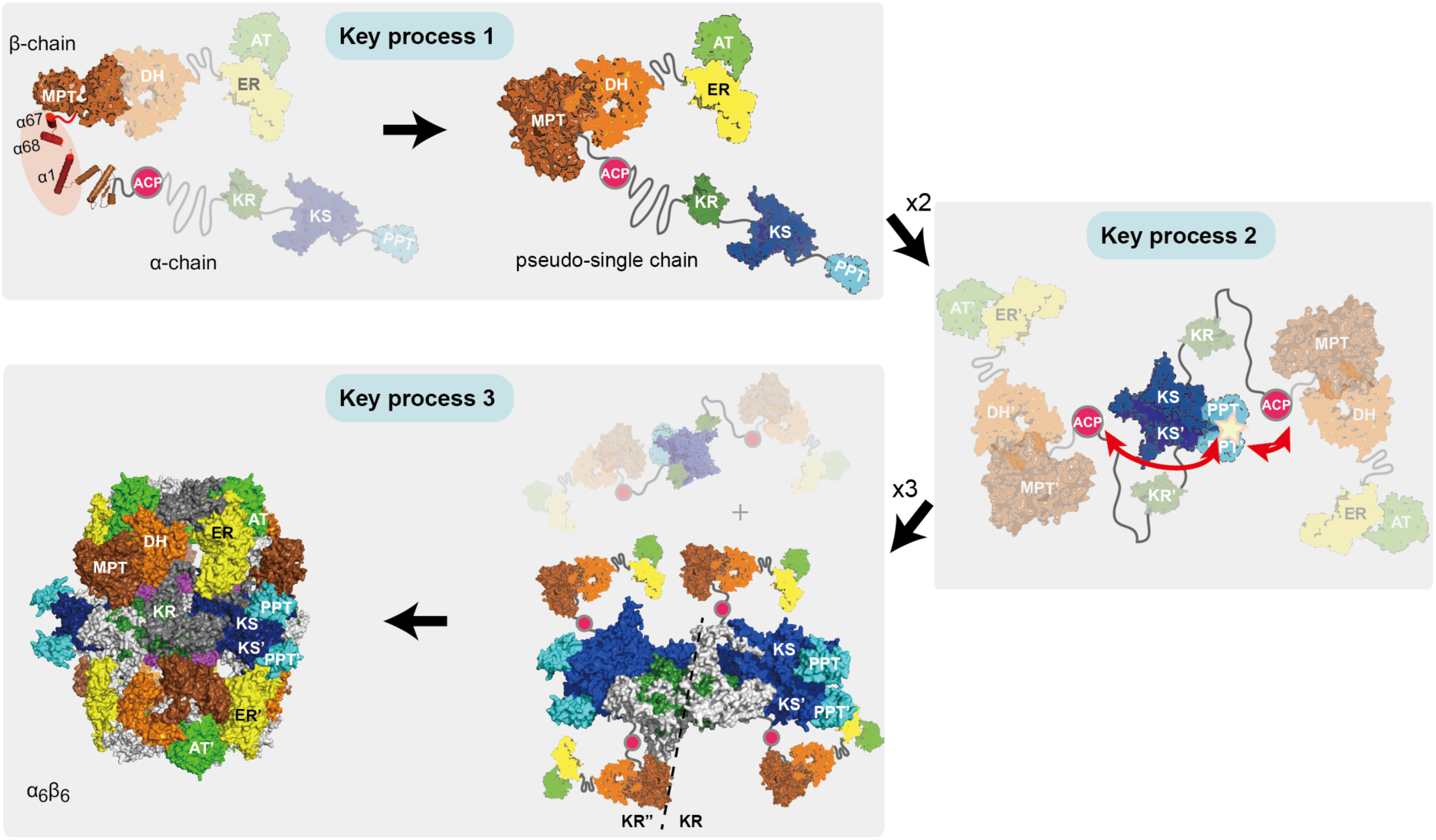
Model for the assembly of *S. cerevisiae* FAS in three key processes. The integration of termini for the formation of the MPT proceeds early in assembly. The KS dimerization, that occurs subsequently or concerted to process 1, establishes dimeric units that act as platform for the phosphopantetheinylation of ACP by PPT. Finally, abstracted in key process 3, the C2 symmetric dimeric units trimerize to overall D3 symmetric barrel-shaped structures.

As shown for *S. cerevisiae* FAS, fFAS assembly is robust and tolerates gene fusion and alternative gene fission (see **Figure S3**). Our data suggest that the key players driving assembly are mainly the catalytic domains that successively interact during assembly. The insertion elements stabilize the final barrel-shaped structure, and seem to be of minor significance for assembly except evolutionary ancient motifs as e.g. DM3 ^22^. fFAS is the most efficient *de novo* fatty acid (FA) synthesizing protein ^9^. This property makes fFAS an attractive object in the endeavor to achieve microbial production of FA^42^. The barrel-shaped fold has recently also been suggested as scaffold that may more generally be repurposed as microbial compartment for synthetic pathways ^20^. Our data suggest that domains AT, ER and DH may indeed be susceptible to domain swapping to putatively enlarge and modify fFAS by new catalytic functions, while they similar show that engineering strategies have to preserve the domains KS, KR and MPT, owing to their structural tasks and roles in the assembly process. Engineering approaches that employ termini of insertion elements as docking or attachment sites may further allow enlarging the fFAS scaffold with new functions. Here, a core FA synthetizing unit may be decorated with FA modifying catalytic functions.

Data presented in this study are valuable for guiding biotechnological fFAS design: A recent approach has taken into account structural aspects and information towards the biogenesis of fFAS, and successfully added a thioesterase (TE) domain to the *S. cerevisiae* FAS by inserting the TE in the ACP linker sequences as well as at the C-terminus of the α-chain ^13^. An alternative approach was recently performed with the *Ascomycota*-type *Yarrowia lipolytica* FAS. For achieving short chain FA production, the C-terminal part of the split MPT was replaced with a TE domain, which, however, hampers the assembly to fully active intact protein, since interfering in pseudo-single chain formation. Such strategies can now be avoided when following the here presented guidelines ^43^.

## Conclusion

The 2.6 MDa fungal fatty acid synthase (fFAS) is among the most elaborate protein complexes known to date, and an interesting object for studying the assembly of multidomain proteins. The fFAS protein family comprises a heterogeneous class of proteins: fFAS are built from one or two polypeptides, and insertion elements that scaffold the barrel structure are non-uniformly distributed among the variants in the fFAS ^2,3^. A conserved assembly pathway needs to reconcile all the individual topologies found in the fFAS family ^26^.

Our approach of analyzing the molecular mechanisms in fFAS assembly was strongly guided by correlating the assembly pathway to the evolutionary development of the fFAS protein family. To disclose the assembly pathway and its molecular mechanisms, we performed *in vivo* mutational studies as well as the *in vitro* analysis of full-length and truncated proteins. We finally show data that characterize fFAS assembly as progressing through three key processes. In the initial step, the polypeptide chains of two-gene encoding fFAS interact via a small interface for pseudo-single chain formation, while the following steps include the formation of larger domain-domain interfaces. We show that the initial interaction is sensitive to small perturbations, which may be exploited in ultra-selective inhibition of *de novo* FA synthesis in antifungal therapy. As one of the most intriguing aspects of this study, the assembly pathway appears to be entirely sequence coded. The assembly does not require external factors nor does it involve stalled intermediate state, i.e. for proof-reading crucial post-translational phosphopantetheinylation. Overall, assembly pathway shows a high plasticity that well corresponds to the heterogeneity of the protein family. This property makes fFAS a suitable scaffold for engineering compartmentalized biosynthetic pathways ^11,13,20^.

## Experimental Procedures

Please find a detailed description of the experimental design in **Supplemental Experimental Procedures**.

### Plasmids and transformation

Yeast plasmids have a pRS backbone with centromeric replication site ^23^ and were cloned with homologous recombination in *S. cerevisiae* or with the Infusion HD cloning kit (Clontech, USA) in *E. coli*. All *FAS1* and *FAS2* derived constructs carry the native promoter and terminator sequences ^24^. Yeast transformation was done with the LiOAc-method ^44^. *E. coli* plasmids have a pET22b backbone (Novagen, USA) and were cloned with the Infusion HD cloning kit (Clontech, USA) (see **Table S1**).

### Creation of FAS deficient *S. cerevisiae* strain *BY.PK1238_KO*

Strains *Y25032* and *Y21061* were transformed with pMF001 and, after two rounds of sporulation, yielded the haploid *Δfas1; Δfas1* strain. Rejection of the rescue plasmid pMF001 was achieved via selection with 5-fluoroorotic acid.

### Protein purification

Wild type FAS from *S. cerevisiae* as well as ΔTM, ΔDM2 and ΔTM ΔDM2 deletion mutants were isolated as Strep-I-tagged proteins from *S. cerevisiae* with basal expression. Purification of other FAS constructs for *in vitro* studies was achieved from homologous expressions in *E. coli*

### Liquid culture growth assay

Cells from single yeast colonies were picked to inoculate 5 mL cultures in appropriate selection medium containing 200 µg/mL geneticin disulfate, free FA (myristic, palmitic and stearic acid, each 50 mg/L) and 1% Tergitol NP-40. After growth at 30 °C and 200 rpm, pre-cultures of same media were inoculated, and grown at 30 °C and 200 rpm to OD(600) 1-14. For 5 mL main cultures in YPD (containing 1% Tergitol NP-40, varying FA concentrations and 200 µg/mL geneticin disulfate), reproducible inocula were obtained by using a standardized inoculum procedure to yield a constant starting OD(600) of 32 × 10^−3^. The cultures were incubated for 24 h at 30 °C and 200 rpm.

### Serial dilution growth assay

Cells were precultured as mentioned above and in 4-fold 1:10 serial dilutions starting from OD(600) 1 transferred onto YPD agar plates without FA. Growth differences were recorded following incubation of the plates for 2-3 days at 30 °C.

### Native PAGE with Western Blot analysis

*S. cerevisiae* cultures were grown to OD(600) 1 to 2 in appropriate selection medium containing 200 µg/mL geneticin disulfate, free FA (myristic, palmitic and stearic acid, each 50 mg/L) and 1% Tergitol NP-40. Cells were lysed with Zymolyase and lysates were concentrated to total protein concentrations between 1 mg/mL and 5 mg/mL. Native-PAGE (3-12% Bis-Tris gels, Novex, Life Technologies, US) was performed with varying volumes to achieve identical total protein amounts for every sample. As reference, a total amount of 0.1 to 0.2 µg purified *S. cerevisiae* FAS was loaded. After electrophoresis in Blue Native buffer system (Serva Electrophoresis GmbH, Germany) and blotting onto a polyvinylidene difluoride membrane (Immobilon-FL, Merck Millipore, Germany) by electro-transfer, FAS proteins were detected with rabbit anti-FAS antiserum ^25^ and horseradish peroxidase conjugated goat anti-rabbit IgG (Pierce, Thermo Fisher Scientific, USA). Luminescence was induced with peroxidase substrate (Carl Roth GmbH, Germany).

### Protein purification and protein biochemical assays

methods for purification of different FAS constructs and fragments as well as for thermal shift and activity assay are given in the supplementary materials and methods.

### CD-spectroscopy

The peptides α1 (MKPEVEQELAHILLTELLAYQ-NH_2_), α1_K2S/E8R (MSPEVEQRLAHILLTELLAYQ-NH_2_) and α67/68 (Ac-VTKEYFQDVYDLTGSEPIKEIIDNWEKYEQ) (CASLO ApS, Denmark) were measured at 40 µM in buffer (100 mM NaPi, pH 7.2) with varying volume fractions of 2,2,2-Trifluoroethanol (Alfa Aesar, Johnson Matthey GmbH, Germany) on a Jasco J810 spectrometer (Jasco GmbH, Germany).

## Acknowledgements

We are grateful to Patrik Johansson for starting the project with us. We thank Michael Thumm (Georg August University Göttingen) for providing anti-FAS antiserum, and the EMBL Heidelberg for providing the plasmid pET28M-Sumo1. We received assistance from Merle Hantsche in performing ACP-PPT interactions studies, and Jurema Schmidt and Veysel Erdel in setting up the Native-PAGE with Western Blotting. We are also grateful to Rudolf Glockshuber, Michael Groll, Werner Kühlbrandt, Rolf Marschalek, Nina Morgner and Harald Schwalbe for valuable discussions on this topic. Furthermore we, again, thank Harald Schwalbe and Harald Hofbauer for support in CD-spectroscopy. This work was supported by a Lichtenberg Grant of the Volkswagen Foundation to M.G. (grant number 85 701).

## Author Contributions

M.F. rationally engineered FAS constructs, set-up and performed the complementation assay, purified FAS, performed CD-spectroscopic studies, analyzed data and wrote the manuscript; B.M. and R.V. purified and analyzed FAS and FAS constructs from recombinant expressions in *E. coli*; M.J. purified FAS constructs from yeast expression, performed activity assays, thermo shift experiments and analyzed the data; K.K. cloned the α_ΔMPT-tev construct and was involved in cloning and expressing of other FAS constructs; P.K. was main responsible in constructing the FAS deficient *S. cerevisiae* strain *BY.PK1238_KO*; L.C. and J.V. performed negative stain electron microscopic studies on the *S. cerevisiae* FAS and the β-chain α-chain fusion construct recombinantly expressed in *E. coli*; D.O. analyzed data; M.G. expressed and analyzed proteins, analyzed data, designed research and wrote the paper.

## Abbreviations

FA: fatty acids
FAS: fatty acid synthase
fFAS: fungal fatty acid synthase
PPT: phosphopantetheine transferase
ACP: acyl carrier protein
KS: ketoacyl synthase
KR: ketoacyl reductase
PKS: polyketide synthase

## References

1. White, S. W., Zheng, J., Zhang, Y.-M. & Rock, C. O. The structural biology of type II fatty acid biosynthesis. Annu Rev Biochem. 74, 791–831 (2005).

2. Maier, T., Leibundgut, M., Boehringer, D. & Ban, N. Structure and function of eukaryotic fatty acid synthases. Q Rev Biophys. 43, 373–422 (2010).

3. Grininger, M. Perspectives on the evolution, assembly and conformational dynamics of fatty acid synthase type I (FAS I) systems. Curr Opin Struct Biol. 25, 49–56 (2014).

4. Leibundgut, M., Jenni, S., Frick, C. & Ban, N. Structural basis for substrate delivery by acyl carrier protein in the yeast fatty acid synthase. Science 316, 288–290 (2007).

5. Lomakin, I. B., Xiong, Y. & Steitz, T. A. The crystal structure of yeast fatty acid synthase, a cellular machine with eight active sites working together. Cell 129, 319–332 (2007).

6. Johansson, P. et al. Inhibition of the fungal fatty acid synthase type I multienzyme complex. Proc Natl Acad Sci U S A. 105, 12803–12808 (2008).

7. Jenni, S. et al. Structure of fungal fatty acid synthase and implications for iterative substrate shuttling. Science 316, 254–261 (2007).

8. Gipson, P. et al. Direct structural insight into the substrate-shuttling mechanism of yeast fatty acid synthase by electron cryomicroscopy. Proc Natl Acad Sci U S A. 107, 9164–9169 (2010).

9. Fischer, M. & Grininger, M. Strategies in megasynthase engineering – fatty acid synthases (FAS) as model proteins. Beilstein J Org Chem. 13, 1204–1211 (2017).

10. Blazeck, J. et al. Harnessing Yarrowia lipolytica lipogenesis to create a platform for lipid and biofuel production. Nat Commun. 5, 3131, (2014).

11. Gajewski, J. et al. Engineering fatty acid synthases for directed polyketide production. Nat Chem Biol. 13, 363–365 (2017).

12. Gajewski, J., Pavlovic, R., Fischer, M., Boles, E. & Grininger, M. Engineering fungal de novo fatty acid synthesis for short chain fatty acid production. Nat Commun. 8, 14650 (2017).

13. Zhu, Z. et al. Expanding the product portfolio of fungal type I fatty acid synthases. Nat Chem Biol. 13, 360–362 (2017).

14. Fischer, M. et al. Cryo-EM structure of fatty acid synthase (FAS) from Rhodosporidium toruloides provides insights into the evolutionary development of fungal FAS. Protein Sci. 24, 987–995 (2015).

15. Enderle, M., McCarthy, A., Paithankar, K. S. & Grininger, M. Crystallization and X-ray diffraction studies of a complete bacterial fatty-acid synthase type I. Acta Crystallogr F Struct Biol Commun. 71, 1401–1407 (2015).

16. Levy, E. D., Boeri Erba, E., Robinson, C. V. & Teichmann, S. A. Assembly reflects evolution of protein complexes. Nature 453, 1262–1265 (2008).

17. Chayakulkeeree, M., Rude, T. H., Toffaletti, D. L. & Perfect, J. R. Fatty acid synthesis is essential for survival of Cryptococcus neoformans and a potential fungicidal target. Antimicrobial Agents and Chemotherapy 51, 3537–3545 (2007).

18. Nguyen, L. N., Trofa, D. & Nosanchuk, J. D. Fatty acid synthase impacts the pathobiology of Candida parapsilosis in vitro and during mammalian infection. PLoS One 4(12):e8421 (2009).

19. Nguyen, L. N. et al. Inhibition of Candida parapsilosis fatty acid synthase (Fas2) induces mitochondrial cell death in serum. PLoS Pathog. 8, e1002879 (2012).

20. Maier, T. Fatty acid synthases: Re-engineering biofactories. Nat Chem Biol. 13, 344–345 (2017).

21. Johansson, P. et al. Multimeric options for the auto-activation of the Saccharomyces cerevisiae FAS type I megasynthase. Structure 17, 1063–1074 (2009).

22. Bukhari, H. S. T., Jakob, R. P. & Maier, T. Evolutionary Origins of the Multienzyme Architecture of Giant Fungal Fatty Acid Synthase. Structure 22, 1775–1785, (2014).

23. Sikorski, R. S. & Hieter, P. A system of shuttle vectors and yeast host strains designed for efficient manipulation of DNA in Saccharomyces cerevisiae. Genetics 122, 19–27 (1989).

24. Chirala, S. S. Coordinated regulation and inositol-mediated and fatty acid-mediated repression of fatty acid synthase genes in Saccharomyces cerevisiae. Proc Natl Acad Sci U S A. 89, 10232–10236 (1992).

25. Egner, R. et al. Tracing intracellular proteolytic pathways. Proteolysis of fatty acid synthase and other cytoplasmic proteins in the yeast Saccharomyces cerevisiae. J Biol Chem. 268, 27269–27276 (1993).

26. Marsh, J. A. et al. Protein complexes are under evolutionary selection to assemble via ordered pathways. Cell 153, 461–470 (2013).

27. Schüller, H. J., Förtsch, B., Rautenstrauss, B., Wolf, D. H. & Schweizer, E. Differential proteolytic sensitivity of yeast fatty acid synthetase subunits alpha and beta contributing to a balanced ratio of both fatty acid synthetase components. Eur J Biochem. 203, 607–614 (1992).

28. Wenz, P., Schwank, S., Hoja, U. & Schueller, H.-J. A downstream regulatory element located within the coding sequence mediates autoregulated expression of the yeast fatty acid synthase gene FAS2 by the FAS1 gene product. Nucleic Acids Res. 29, 4625–4632 (2001).

29. Buck, M. Trifluoroethanol and colleagues: cosolvents come of age. Recent studies with peptides and proteins. Q Rev Biophys. 31, 297–355 (1998).

30. Sievers, F. et al. Fast, scalable generation of high-quality protein multiple sequence alignments using Clustal Omega. Mol Syst Biol. 7, 1–6 (2011).

31. Buchan, D. W. A., Minneci, F., Nugent, T. C. O., Bryson, K. & Jones, D. T. Scalable web services for the PSIPRED Protein Analysis Workbench Nucleic Acids Res. 41, W340–W348 (2013).

32. Stuible, H. P., Meurer, G. & Schweizer, E. Heterologous expression and biochemical characterization of two functionally different type I fatty acid synthases from Brevibacterium ammoniagenes. Eur J Biochem. 247, 268–273 (1997).

33. Ciccarelli, L. et al. Structure and conformational variability of the Mycobacterium tuberculosis fatty acid synthase multienzyme complex. Structure 21, 1251–1257 (2013).

34. Lambalot, R. H. et al. A new enzyme superfamily - the phosphopantetheinyl transferases. Chem Biol. 3, 923–936 (1996).

35. Fichtlscherer, F., Wellein, C., Mittag, M. & Schweizer, E. A novel function of yeast fatty acid synthase. Subunit alpha is capable of self-pantetheinylation. Eur J Biochem. 267, 2666–2671 (2000).

36. Fisher, M. C. et al. Emerging fungal threats to animal, plant and ecosystem health. Nature 484, 186–194 (2012).

37. Shapiro, R. S., Robbins, N. & Cowen, L. E. Regulatory circuitry governing fungal development, drug resistance, and disease. Microbiol Mol Biol Rev. 75, 213–267 (2011).

38. Chayakulkeeree, M. & Perfect, J. R. Cryptococcosis. Infect Dis Clin North Am. 20, 507–544 (2006).

39. Pfaller, M. A. & Diekema, D. J. Epidemiology of invasive candidiasis: a persistent public health problem. Clin Microbiol Rev. 20, 133–163 (2007).

40. Uversky, V. N. The most important thing is the tail: multitudinous functionalities of intrinsically disordered protein termini. FEBS Lett 587, 1891-1901(2013).

41. Kikuchi, S., Rainwater, D. L. & Kolattukudy, P. E. Purification and characterization of an unusually large fatty acid synthase from Mycobacterium tuberculosis var. bovis BCG. Arch Biochem Biophys. 295, 318–326 (1992).

42. d’Espaux, L., Mendez-Perez, D., Li, R. & Keasling, J. D. Synthetic biology for microbial production of lipid-based biofuels. Curr Opin Chem Biol. 29, 58–65 (2015).

43. Xu, P., Qiao, K., Ahn, W. S. & Stephanopoulos, G. Engineering Yarrowia lipolytica as a platform for synthesis of drop-in transportation fuels and oleochemicals. Proc Natl Acad Sci U S A. 113, 10848–10853 (2016).

44. Gietz, R. D. & Schiestl, R. H. High-efficiency yeast transformation using the LiAc/SS carrier DNA/PEG method. Nat Protoc. 2, 31–34 (2007).

